# A Tn-seq screen of *Streptococcus pneumoniae* uncovers DNA repair as the major pathway for desiccation tolerance and transmission

**DOI:** 10.1101/2020.11.09.375980

**Authors:** Allison J. Matthews, Hannah M. Rowe, Jason W. Rosch, Andrew Camilli

## Abstract

*Streptococcus pneumoniae* is an opportunistic pathogen that is a common cause of serious invasive diseases such as pneumonia, bacteremia, meningitis, and otitis media. Transmission of this bacterium has classically been thought to occur through inhalation of respiratory droplets and direct contact with nasal secretions. However, the demonstration that *S. pneumoniae* is desiccation tolerant, and therefore environmentally stable for extended periods of time, opens up the possibility that this pathogen is also transmitted via contaminated surfaces (fomites). To better understand the molecular mechanisms that enable *S. pneumoniae* to survive periods of desiccation, we performed a high throughput transposon sequencing (Tn-seq) screen in search of genetic determinants of desiccation tolerance. We identified 42 genes whose disruption reduced desiccation tolerance, and 45 genes that enhanced desiccation tolerance. The nucleotide excision repair pathway was the most enriched category in our Tn-seq results, and we found that additional DNA repair pathways are required for desiccation tolerance, demonstrating the importance of maintaining genome integrity after desiccation. Deletion of the nucleotide excision repair gene *uvrA* resulted in decreased transmission efficiency between infant mice, indicating a correlation between desiccation tolerance and pneumococcal transmission. Understanding the molecular mechanisms that enable pneumococcal persistence in the environment may enable targeting of these pathways to prevent fomite transmission, thereby preventing the establishment of new colonization and any resulting invasive disease.

## INTRODUCTION

For pathogens with no environmental reservoir, transmission between hosts is necessary for the species to survive. In the case of *Streptococcus pneumoniae* (the Pneumococcus), a common member of the human nasopharyngeal microbiome, most transmission events result in asymptomatic and transient colonization which has been termed the carrier state (1–4). However, in susceptible individuals such as children and the elderly, *S. pneumoniae* can be aspirated into the lungs resulting in pneumonia and invasive diseases such as bacteremia and meningitis (5). Due to high carriage rates of *S. pneumoniae* within the population, invasive pneumococcal disease continues to be a leading cause of lower respiratory morbidity and mortality as well as a significant socioeconomic burden (6–8). As colonization precedes invasive pneumococcal disease, developing ways to prevent colonization, such as limiting fomite transmission, would serve to reduce the incidence of invasive disease.

The prevailing model of pneumococcal transmission posits that transmission occurs via respiratory droplets and direct contact with nasal secretions. However, previous work has demonstrated that *S. pneumoniae* can survive long periods of desiccation (9, 10). Upon subsequent rehydration, a proportion of the bacteria were found to remain viable and capable of establishing colonization. Thus, environmentally stable bacteria desiccated on surfaces, also referred to as fomites, may serve as an alternate source of pneumococcal infection.

Surfaces contaminated with infectious microbes are an important mode of transmission for a number of pathogens (11–18). In particular, fomites have been demonstrated to be a frequent source of nosocomial infections (19–24). Therefore, the demonstration that *S. pneumoniae* can be isolated from surfaces in a daycare provides evidence that fomite reservoirs of the bacterium exist in the community (10, 25). As *S. pneumoniae* is desiccation tolerant for an extended period of time, it is likely that the bacterium uses fomite transmission as one of multiple strategies to reach new hosts. Furthermore, increased desiccation tolerance of a pyruvate oxidase mutant has been shown to correlate with improved transmission between infant mice in a murine model of pneumococcal transmission, providing support to the hypothesis of pneumococcal fomite transmission (26).

Although fomites may constitute an important mode of transmission for *S. pneumoniae*, little is known about the molecular mechanisms that enable *S. pneumoniae* to remain stable in the environment as the bacteria desiccate and are left without access to nutrients. Desiccation is theorized to impose an enormous amount of stress on an organism. Some of these stresses include DNA damage, protein damage, osmotic shock, oxidative damage, protein denaturation and cross-linking, and reduced membrane fluidity (27). These challenges are so great that the majority of bacteria are unable to survive extended periods of desiccation (28). Therefore, the pneumococcus must have evolved mechanisms to cope with the challenges imposed by desiccation. In this study, we used a high throughput mutant screening approach to identify genes that are involved in the desiccation tolerance response of *S. pneumoniae* in order to better characterize environmental persistence of the bacterium.

## RESULTS

### Tn-seq screen to identify genes involved in *S. pneumoniae* desiccation tolerance

To uncover which *S. pneumoniae* factors are required for desiccation tolerance, we employed a high-throughput transposon sequencing (Tn-seq) approach (29). In vitro transposition of a mini-transposon and subsequent transformation of the transposed DNA into bacteria produced a library of ~64,000 unique insertion mutants in the serotype 2 strain D39. This high-complexity library was then screened for sensitivity to desiccation using a previously described desiccation assay (9). To perform the desiccation, bacteria were grown to near-confluence on blood agar and then collected and spread thinly on polystyrene petri plate lids and left in the dark to desiccate for 48 hours. In order to isolate survivors, desiccated bacteria were resuspended and plated on blood agar and then grown overnight.

To prepare the libraries for sequencing, genomic DNA was isolated from the pooled desiccation survivors as well as the input library. The genomic junctions of all transposon insertion mutants were amplified by HTML-PCR as described (29) and each sample was uniquely barcoded. The location of each transposon insertion was then identified using massively parallel sequencing on the Illumina platform and the relative frequency of each mutant within the library was then determined using normalized read counts. Frequencies of each unique insertion mutant were compared from before and after desiccation and this was used to calculate a fitness (*W*) value for each insertion. Mean fitness of a gene was then calculated by averaging the fitness of all transposon insertions within a gene.

As expected, the majority of genes when disrupted by the transposon had a neutral impact on bacterial fitness during desiccation, resulting in a fitness of ~1 (Fig.1; Supplemental Table S1). All genes showing a 20% or greater change in fitness (*W*) with a P value below 2.33×10^−4^ (−log[Pvalue]>3.633) were considered to have a significant deviation from wild-type. Both genes that contribute to desiccation tolerance (desiccation sensitive) and ones that hindered it (desiccation resistant) were identified (Fig. 1A and B, respectively). Reproducibility was high between the two biological replicates (Pearson’s correlation, R=0.801), providing confidence in the results of the screen (Fig.1C). In total, this screen identified 42 genes whose disruption by transposon insertion render the bacterium desiccation sensitive and 45 genes that resulted in improved survival (Table 1).

**Table 1.** Putative desiccation tolerance genes from the Tn-seq screen. a. Average fitness between the two biological replicates (In cases where the gene did not meet analytical cutoffs for read counts and Tn insertions in one biological replicate, only the fitness of the significant replicate is displayed).

| Desiccation Sensitive |  |  | Desiccation Resistant |  |  |
| --- | --- | --- | --- | --- | --- |
| D39 Locus | Gene | Fitness<sup>a</sup> | D39 Locus | Gene | Fitness<sup>a</sup> |
| SPD_1998 | *adcB* | 0.04 | SPD_0670 |  | 1.21 |
| SPD_1542 | *stkP* | 0.05 | SPD_0700 | *pepN* | 1.21 |
| SPD_1099 |  | 0.06 | SPD_1677 | *rafE* | 1.21 |
| SPD_2000 | *adcR* | 0.09 | SPD_1418 |  | 1.21 |
| SPD_1098 |  | 0.09 | SPD_1491 |  | 1.22 |
| SPD_0185 |  | 0.12 | SPD_0542 | *pepV* | 1.22 |
| SPD_1084 | *vicK* | 0.13 | SPD_1676 | *rafF* | 1.22 |
| SPD_0320 | *cps2T* | 0.14 | SPD_1166 |  | 1.23 |
| SPD_1797 | *ccpA* | 0.17 | SPD_1068 | *udk* | 1.23 |
| SPD_2055 | *guaB* | 0.18 | SPD_0641 | *manA* | 1.23 |
| SPD_1096 | *uvrB* | 0.20 | SPD_0685 | *gor* | 1.24 |
| SPD_0980 | *prs2* | 0.25 | SPD_1667 | *amiF* | 1.24 |
| SPD_1740 | *cinA* | 0.26 | SPD_0820 | *rluD* | 1.24 |
| SPD_0330 | *rfbB* | 0.26 | SPD_1669 | *amiD* | 1.25 |
| SPD_0129 | *gidA* | 0.26 | SPD_1971 |  | 1.25 |
| SPD_0538 | *uvrC* | 0.26 | SPD_1635 | *galR* | 1.25 |
| SPD_1779 |  | 0.31 | SPD_0437 | *ribU* | 1.26 |
| SPD_1295 | *Hemolysin III* | 0.33 | SPD_1409 | *msmK* | 1.26 |
| SPD_0176 | *uvrA* | 0.35 | SPD_1450 | *mntR* | 1.26 |
| SPD_0010 |  | 0.35 | SPD_1668 | *amiE* | 1.26 |
| SPD_2032 | *pde1* | 0.35 | SPD_1670 | *amiC* | 1.26 |
| SPD_1121 |  | 0.38 | SPD_1487 |  | 1.26 |
| SPD_1778 | *rmuC* | 0.39 | SPD_0787 | *pepX* | 1.26 |
| SPD_1549 | *rnY* | 0.40 | SPD_1487 |  | 1.27 |
| SPD_0646 |  | 0.40 | SPD_1634 | *galK* | 1.27 |
| SPD_1104 |  | 0.44 | SPD_1671 | *amiA* | 1.28 |
| SPD_0336 | *pbp1A* | 0.46 | SPD_1633 | *galT-2* | 1.28 |
| SPD_0996 |  | 0.48 | SPD_1165 |  | 1.28 |
| SPD_0268 |  | 0.50 | SPD_1171 |  | 1.28 |
| SPD_1298 |  | 0.55 | SPD_0064 | *cpsR* | 1.30 |
| SPD_1333 |  | 0.56 | SPD_0819 | *lspA* | 1.32 |
| SPD_0342 | *mapZ* | 0.57 | SPD_2046 | *cbiQ* | 1.32 |
| SPD_1821 | *pbp2A* | 0.58 | SPD_0065 | *bgaC* | 1.32 |
| SPD_1622 |  | 0.62 | SPD_2047 | *cbiO1* | 1.33 |
| SPD_0953 | *ppc* | 0.64 | SPD_2048 | *cbiO2* | 1.33 |
| SPD_1867 |  | 0.66 | SPD_1170 | *appA* | 1.40 |
| SPD_2068 | *htrA* | 0.67 | SPD_1375 |  | 1.41 |
| SPD_0366 | *recD2* | 0.69 | SPD_1169 | *appB* | 1.42 |
| SPD_1621 |  | 0.71 | SPD_1167 | *appD* | 1.42 |
| SPD_0068 | *gadE* | 0.71 | SPD_1168 | *appC* | 1.45 |
| SPD_0535 | *murM/fibA* | 0.72 | SPD_0513 | *cysE* | 1.45 |
| SPD_1274 | *guaA* | 0.75 | SPD_1678 | *agaN* | 1.47 |
|  |  |  | SPD_0150 | *gshT* | 1.53 |
|  |  |  | SPD_0818 | *cmbR* | 1.55 |
|  |  |  | SPD_0636 | *spxB* | 1.67 |

**Figure 1.**
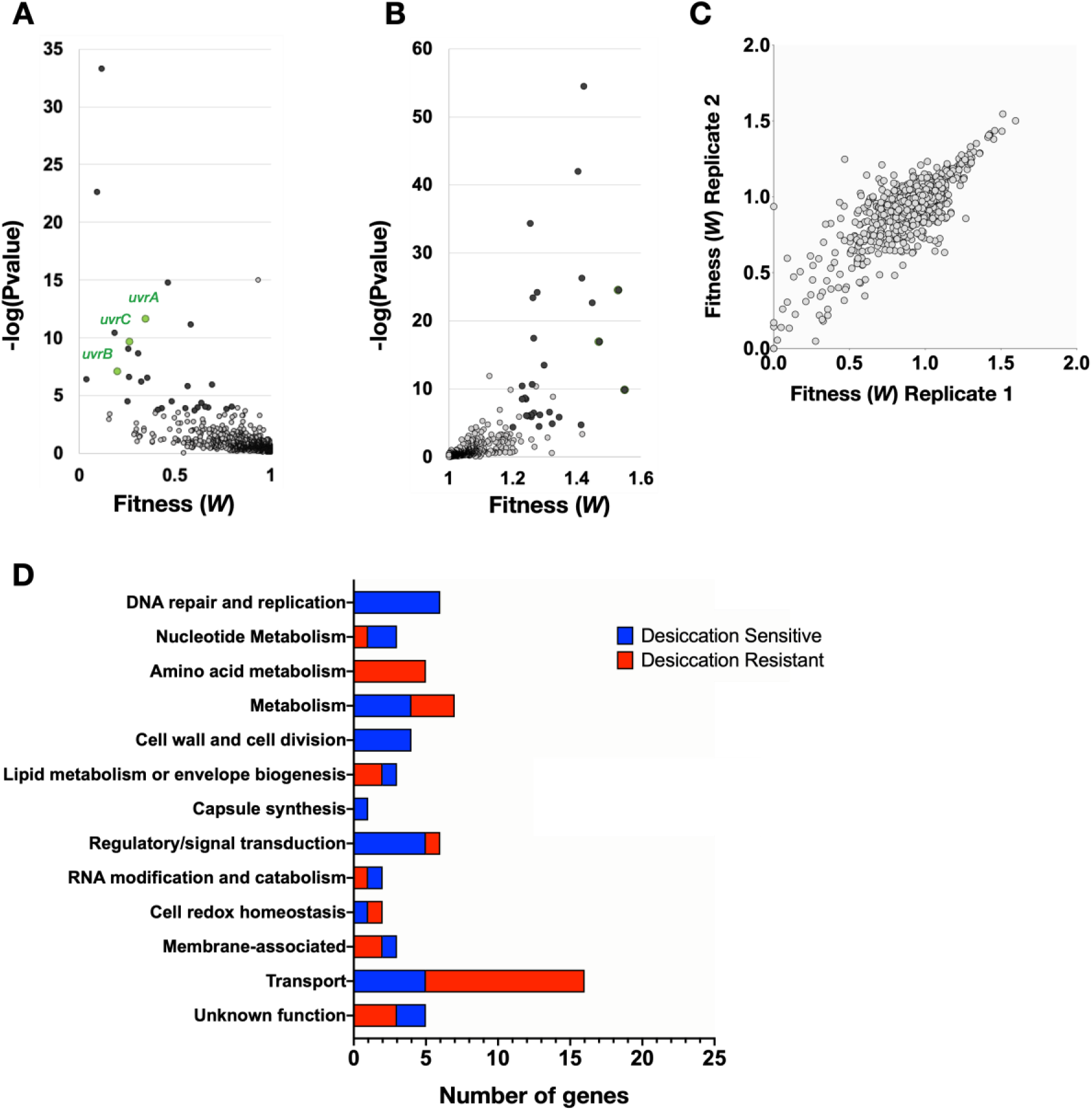
Desiccation tolerance Tn-seq results. Volcano plots of Tn-seq results display statistical significance against fitness for both (A) desiccation sensitive transposon mutants and (B) desiccation resistant mutants. Mutants with a 20% or greater change in fitness that are above the -log(Pvalue) cutoff of 3.633 are highlighted in black. The three components of the highly enriched nucleotide excision repair complex UvrABC are highlighted in green. (C) Reproducibility of the two biological replicates is demonstrated by a Pearson’s correlation of R=0.801. (D) Significant hits from the screen were categorized by function using annotations and GO terms from Kegg genome database and UniProt. The number of genes within each category is quantified on the X-axis.

These genes were categorized by function using annotations and GO terms from Kegg genome database and UniProt (Fig. 1D). Multiple categories of gene disruption rendered the bacteria desiccation sensitive. In particular, genes required for DNA repair and replication (6) and nucleotide metabolism (3) were abundant among the sensitive mutants. To further support the significance of this category of genes, a Gene Ontology (GO) enrichment for cellular components revealed that the excinuclease repair complex UvrABC, which carries out nucleotide excision repair, was enriched 23-fold among our hits. This emphasizes the importance of repairing DNA damage after desiccation and suggests that there is substantial DNA damage that occurs. This is well supported by work in other bacteria that demonstrates the necessity of DNA repair for successful desiccation resistance (30–32). Other functional categories that render the bacterium sensitive to desiccation pertain to composition of the membrane and cell wall. These include the penicillin binding proteins *pbp1A* and *pbp2A* which are responsible for modifying the cell wall, as well as, cardiolipin synthetase which produces the lipid cardiolipin that increases membrane fluidity. Functional categories that result in desiccation resistance include 12 metabolic genes of which five are involved in amino acid metabolism, and 16 different transporters, 3 of which encode sugar transporters. These categories of gene indicate a role for metabolism in desiccation tolerance.

### Validation of putative desiccation tolerance genes

In order to validate the results of our screen, we used allelic replacement to produced deletion mutants of 28 genes. Genes were selected for validation if they had a substantial fitness change, are not pleiotropic in other conditions (33). Genes of known and unknown function were chosen. These deletion mutants were then tested in desiccation tolerance competitions with wild-type. Each mutant was mixed at a 1:1 ratio with wild-type and then plated for overnight growth. The plate grown bacteria were then challenged with a 4 day desiccation and a competitive index (CI) was calculated as the ratio of mutant/wild type in the output divided by the ratio from the input. Similar to the fitness values, a CI less than 1 represents a defect in desiccation tolerance, while a CI greater than 1 represents improved survival. We found that 22 genes validated with competitive indices that were significantly different than that of a neutral gene deletion, *SPD_0022* (Fig. 2; Supplemental Table S2). The majority of genes validated in desiccation competition assays demonstrating the robustness of our screen.

**Figure 2.**
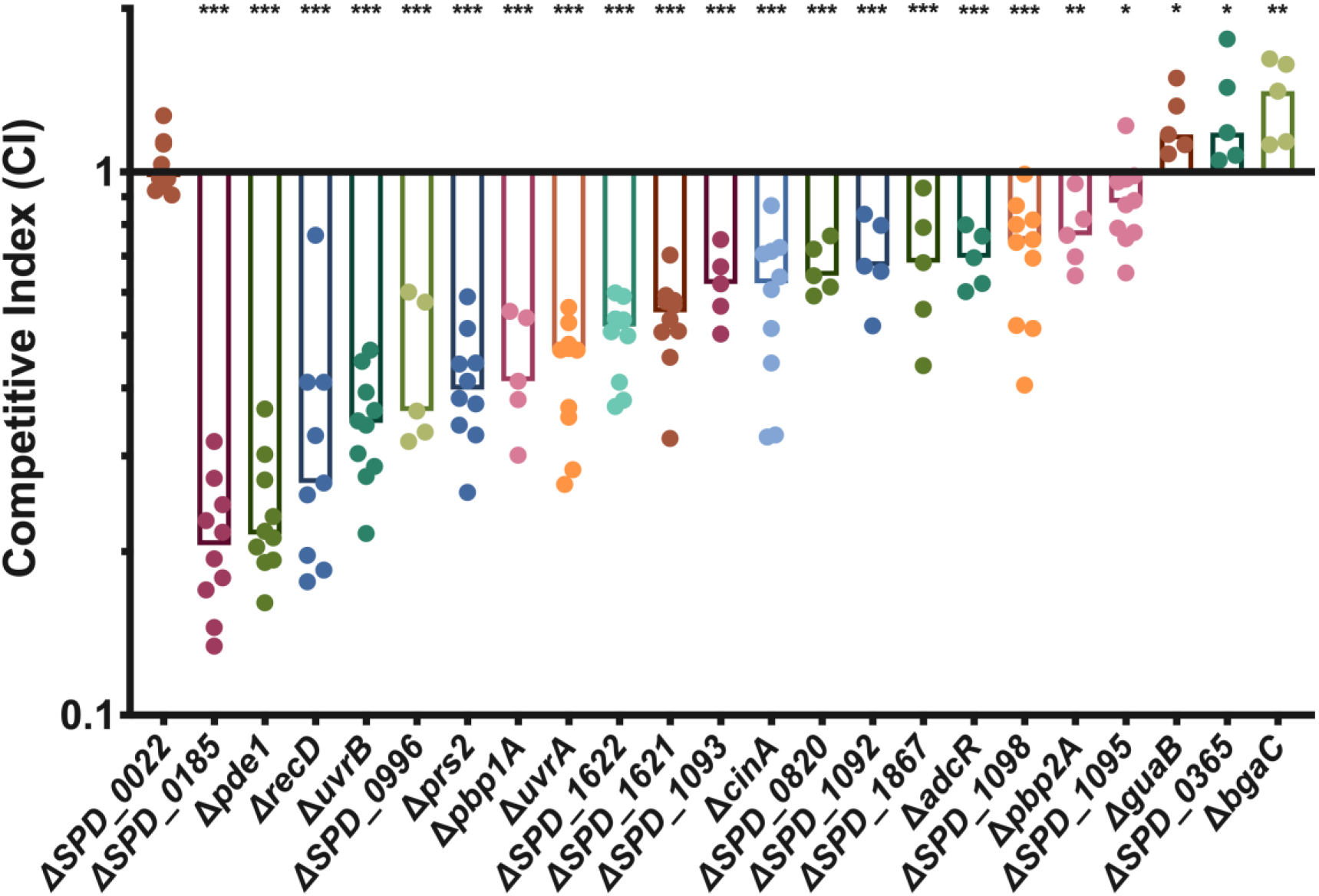
Competitive indices of desiccation tolerance genes. 28 putative desiccation tolerance genes identified in the Tn-seq were deleted and then tested in a 4-day desiccation competition assay against wild-type. Strains in this figure are the 22 deletion mutants that validated in addition to a neutral gene *SPD_0022*. Competitive index was calculated as the ratio of mutant to wild-type after desiccation divided by the ratio before desiccation. The median for each mutant is represented with a bar. Statistical analyses were performed using a non-parametric Mann Whitney U two sample rank test comparing each mutant against the neutral gene SPD_0022 (***, P≤0.001; **, P≤0.002; *, P≤0.033).

We found that both nucleotide excision repair genes tested, *uvrA* and *uvrB*, had a significant defect in desiccation tolerance resulting in median competitive indices of 0.47 and 0.35 (Fig. 2). In addition, the homologous recombination helicase *recD* and nucleotide biosynthetic gene *prs2* also displayed significant fitness defects. Due to the significant enrichment of the excinuclease DNA repair complex and the validation of other DNA repair and maintenance genes we chose to further characterize the impact of DNA repair on desiccation tolerance.

### DNA repair pathways involved desiccation tolerance

Previous work has demonstrated that the drying of bacteria results in extensive DNA damage (34–36). However, the types of DNA damage that occur in desiccating bacteria have been theorized, but there is little direct evidence. To genetically dissect the specific types of DNA damage that are occurring during desiccation, we chose to delete a variety of DNA repair genes, including some not identified in our Tn-seq screen because they were above the P value cutoff. Because specific DNA repair pathways are required for resolving particular DNA lesions, increased desiccation sensitivity resulting from disruption of a DNA repair pathway would suggest a particular type of damage is occurring.

Due to the general essentially of DNA repair for bacterial viability, deletion of many DNA repair genes is lethal. For this reason, we selected genes that function in specific DNA repair pathways but are not essential. We tested multiple genes in the nucleotide excision repair (NER) pathway, including two that are part of the core NER complex (*uvrA, uvrB*) as well as a gene that is only involved in transcription coupled NER (*MFD*). Deletion of *uvrA* and *uvrB* resulted in a significant competitive disadvantage in desiccation survival, while deletion of *MFD* had a neutral effect on desiccation tolerance (Fig. 3). This suggests that the global genome repair pathway of NER is important for desiccation tolerance, but transcription coupled repair is dispensable. We were able to complement *uvrA* at a neutral locus in the chromosome, demonstrating that the *uvrA* deletion was indeed responsible for the observed desiccation sensitivity (Fig 3). To query the significance of homologous recombination (HR), we deleted the HR helicase *recD* and found that this results in a significant loss of viability after desiccation, suggesting homologous recombination is necessary for desiccation tolerance. Next we deleted two glycosylases (*mutM*, *mutY*) involved in base excision repair (BER) and found that both glycosylases have a competitive disadvantage, although the competitive index of Δ*mutM* is significantly lower than that of Δ*mutY*, suggesting that it has a greater impact on repairing DNA damage resulting from desiccation (Fig. 3). Finally, we tested three factors involved in mismatch repair (MMR) (*xseA*, *mutL*, *mutS1*). MutS1 and MutL act in a stepwise fashion with MutS first recognizing the nucleotide mismatch followed by binding of MutL which will recruit an endonuclease to the complex. Neither of these genes displayed a competitive disadvantage in desiccation, suggesting that mismatches are not the primary type of DNA damage occurring during desiccation (Fig. 3). The desiccation sensitivity displayed by Δ*xseA* (Fig. 3), a bi-directional single-stranded DNA exonuclease (ExoVII) that hydrolyzes single stranded DNA can be explained by the fact that this protein is involved in three different DNA repair pathways: mismatch repair, single strand break repair, and homologous recombination. Based on the neutral impact of *mutL* and *mutS* deletion, we suggest that XseA is likely required for repairing single and double strand breaks after desiccation, and not mismatched nucleotides. This makes sense as a desiccated bacterium is likely dormant and not actively replicating its genome, which is where replication errors usually occur.

**Figure 3.**
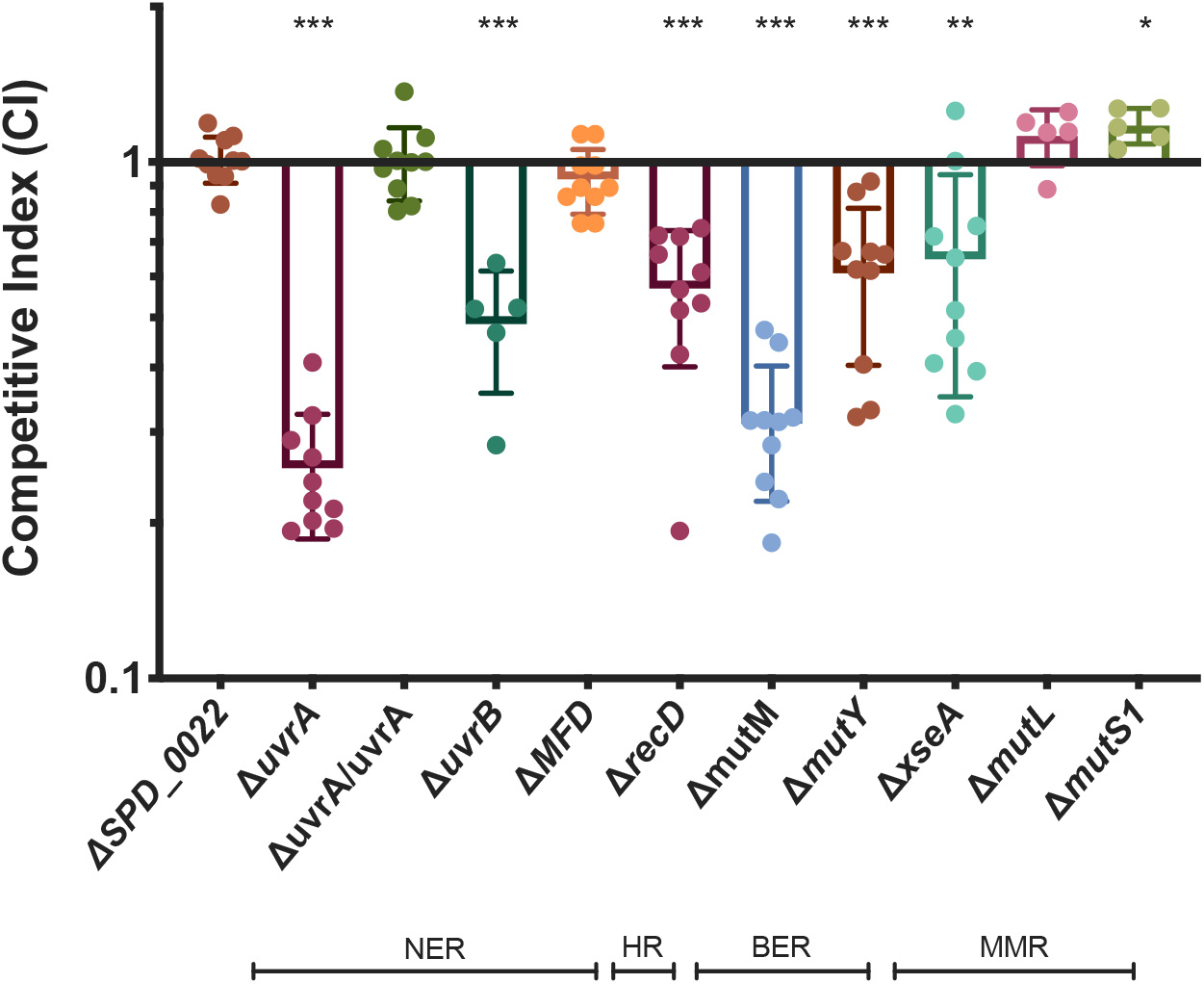
Multiple DNA repair pathways are required for desiccation tolerance. 4-day desiccations were performed on mutants representing a number of DNA repair pathways: Nucleotide excision repair (NER), homologous recombination (HR), base excision repair (BER), and mismatch repair (MMR). Δ*uvrA*/*uvrA* is the *uvrA* deletion mutant with the full gene and native promoter complemented on the chromosome at neutral gene locus *SPD_0022*. Competitive index was calculated as the ratio of mutant to wild-type after desiccation over the input ratio. Statistical analyses were performed using a non-parametric Mann Whitney U two sample rank test comparing each mutant against the neutral gene SPD_0022 (***, P≤0.001; **, P≤0.002; *, P≤0.033).

Having identified BER pathway genes *mutM* and *mutY*, which have both been characterized to repair oxidatively damaged guanines (8-oxoG) (37), we wanted to see if endogenous hydrogen peroxide production was responsible for oxidative damage that may be repaired by BER. *S. pneumoniae* is well known to produce hydrogen peroxide without a detoxification mechanism. The primary producer of hydrogen peroxide is pyruvate oxidase (SpxB) (38), which when deleted resulted in improved desiccation resistance in our screen (Table 1). This desiccation resistance was recapitulated in competition against wild-type (Fig. 4). To probe the impact of hydrogen peroxide production on DNA damage during desiccation, we performed desiccation competitions where we removed the majority of hydrogen peroxide from the system by deleting *spxB* in both the wild-type background and our DNA repair mutants. If the DNA repair mutant were responsible for repairing oxidative damage to the DNA caused by endogenous hydrogen peroxide, we would expect to see an abrogation of the fitness defect when *spxB* is deleted. Deletion of *spxB* caused a slight increase in competitive index of the Δ*mutM* or Δ*uvrA* mutants but the differences were not significant (Fig. 4), suggesting that endogenous hydrogen peroxide production by SpxB is not responsible for the majority of DNA damage that is repaired by either of these DNA repair pathways. This suggests that the improved desiccation tolerance of Δ*spxB* may have more to do with the metabolic role of SpxB in carbon utilization as opposed to its production of hydrogen peroxide as a metabolic byproduct.

**Figure 4.**
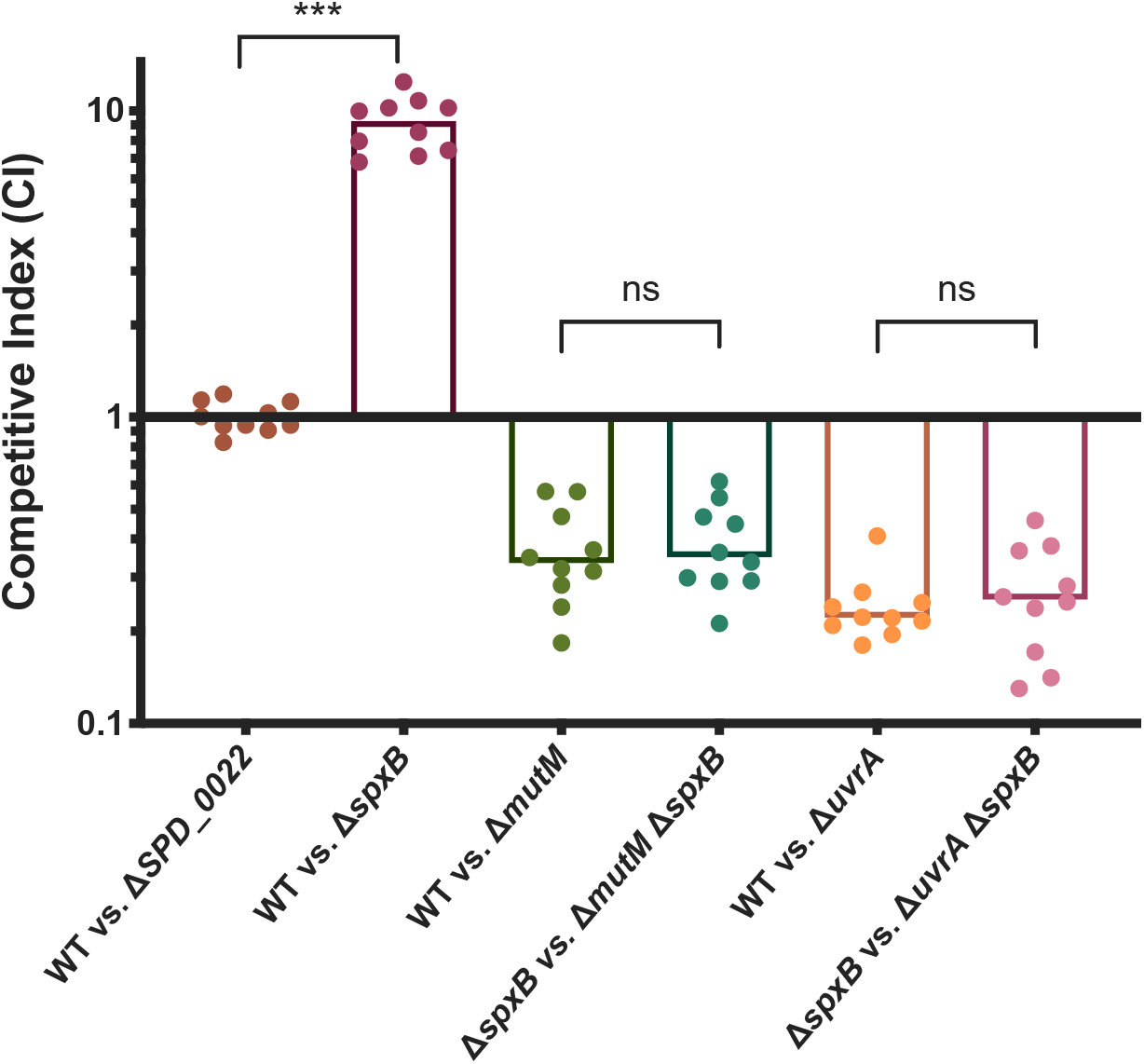
Hydrogen peroxide produced by SpxB is not a primary cause of DNA damage during desiccation. Pyruvate oxidase (SpxB) is responsible for the majority of hydrogen peroxide produced by *S. pneumoniae*. In order to determine if endogenous hydrogen peroxide results in oxidative DNA damage that is repaired by MutM or UvrA, we deleted *spxB* both the wild-type and DNA repair mutant backgrounds. Competitive indices were calculated as the ratio of mutant to wild-type (or double mutant to single mutant) after desiccation compared to the input. Statistical analyses were performed using a non-parametric Mann Whitney U two sample rank test (***, P≤0.001, ns = non-significant).

In order to confirm that the UvrABC complex performs a similar function to its well characterized homolog in *Escherichia coli*, we challenged the Δ*uvrA* mutant with UV irradiation. A deletion in any one of the three components of the NER complex should successfully abrogate its function as all three are required to make a functional complex (39). The *uvrA* deletion mutant was significantly more susceptible to UV treatment, resulting in a 3-log reduction in survival below that of wild-type (Fig. 5). We were able to rescue this phenotype by complementing the *uvrA* gene back at a neutral gene locus, resulting in wild-type survival (Fig. 5). Having confirmed that *uvrA* has a significant impact on desiccation survival and that it’s behavior mimics that of its homologs in other bacteria, we wanted to investigate the impact of *uvrA* deletion and other desiccation mutants on transmission.

**Figure 5.**
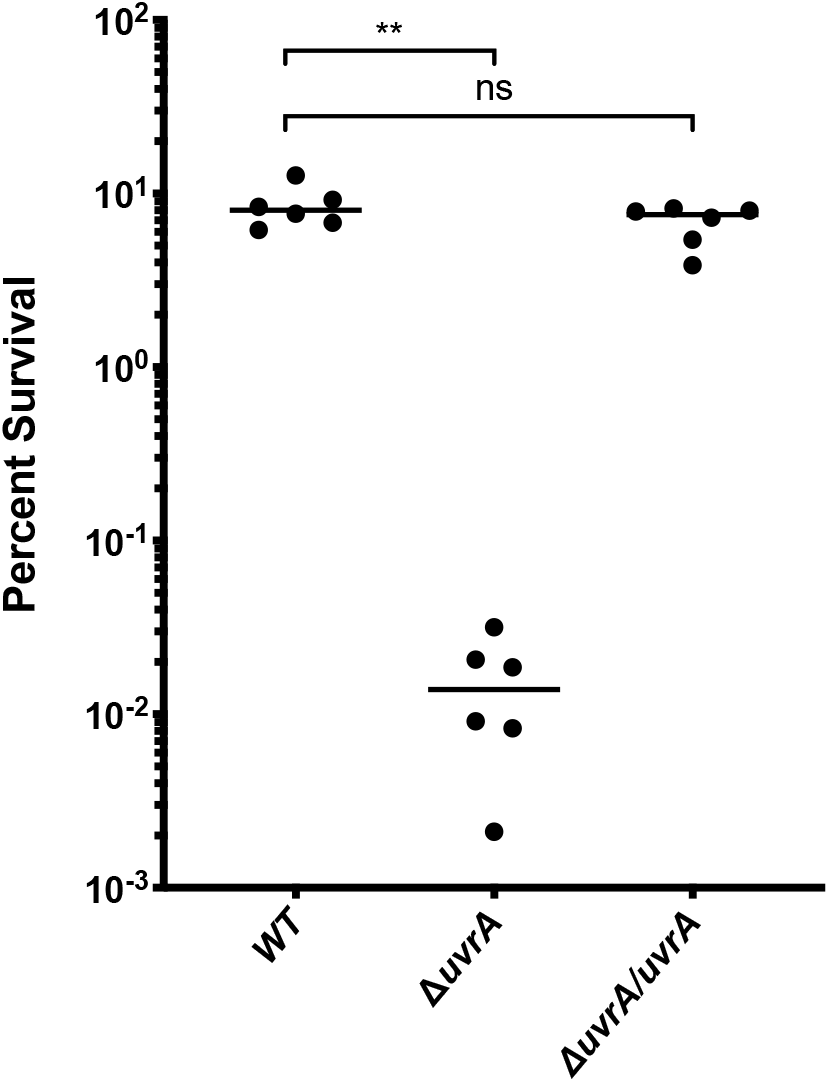
Bacterial survival after UV irradiation. Exponentially growing cultures of *S. pneumoniae* strains were washed and resuspended in PBS and then challenged with 15 millijoules of ultraviolet (UV) light. Percent survival was quantified by plating bacteria for CFU before and after UV exposure. The *uvrA* deletion mutant (Δ*uvrA*) was complemented (Δ*uvrA*/*uvrA*) by placing the full gene and native promoter at neutral gene locus *SPD_0022*. Statistical analyses were performed using a non-parametric Mann-Whitney U two sample rank test (**, P≤0.002, ns = non-significant).

### Transmission efficiency of selected desiccation mutants

As we hypothesize that fomite transmission of *S. pneumoniae* is likely an important method of reaching new hosts, we wanted to see if our desiccation tolerance mutants would impact how efficiently bacteria are passed between mice in a murine model of transmission. Previous work has shown a correlation between transmission efficiency and desiccation tolerance using Δ*spxB;* spxB deletion results in both improved desiccation tolerance as well as increased transmission efficiency (26). Four hits from our screen were selected to be tested in the transmission assay: *uvrA*, *bgaC*, *SPD_1622*, and *SPD_0996*.

The transmission assay was performed by colonizing half of a mouse litter with serotype 19F *S. pneumoniae* (BHN97). These colonized mice are the pneumococcal donors, while the uncolonized littermates are the contact mice. All mice were then returned to their cage with the dam and transmission was tracked over the next 10 days by tapping the nares of the mice against a plate. Detection of colonies on two subsequent days was considered a colonization event. We found that transmission efficiency of the Δ*uvrA* mutant was significantly reduced as compared to wild-type (BHN97) (Fig. 6). The decreased transmission rate of Δ*uvrA* is not due to lower levels of colonization from the donor mice as colonization levels were assessed at the end of the experiment and there was no significant difference between wild-type and Δ*uvrA*. There was no significant difference in transmission efficiency in the other mutants tested, except for *bgaC* which also had reduced levels of colonization in the donor mice (Fig. S1). Altogether, this demonstrates a correlation between decreased desiccation tolerance and lower transmission efficiency.

**Figure 6.**
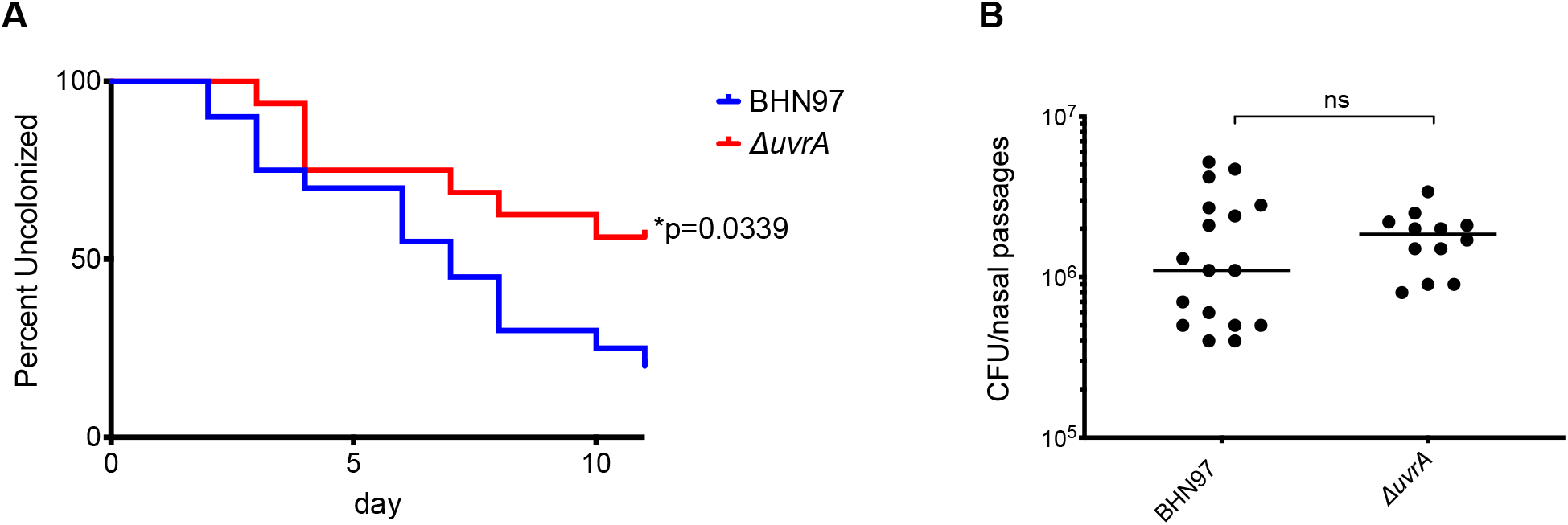
Transmission efficiency of a *uvrA* mutant. Litters of 4-day old, C57/BL6 mice were split into two groups. The first group was colonized with a wild-type or mutant strain of serotype 19F *S. pneumoniae*, while the second half was left uncolonized. All mice were then returned to the cage and uncolonized mice were surveyed daily for transmission by tapping the nares of each mouse against a blood agar plate. A colonization event was defined as detectable CFU on two subsequent days. (A) Transmission of wild-type (BHN97) and Δ*uvrA* was tracked over the course of 10 days. (B) Colonization levels of all donor mice were assessed at the end of the experiment. Statistics were performed with Mantel-cox log-rank test for the transmission assay and Mann-Whitney U two sample rank test for the colonization (*, P≤0.033; ns = non-significant).

## DISCUSSION

*S. pneumoniae* has been demonstrated to be desiccation tolerant, surviving in a dehydrated state for up to 30 days (9, 10). However, little is known about the mechanisms that enable the bacterium to persist in this state. Here we have used transposon insertion sequencing (Tn-seq) to investigate the genetic factors that influence desiccation tolerance of *S. pneumoniae*. We screened approximately 64,000 unique transposon insertion mutants using a 2-day desiccation assay on a plastic surface. After stringent analysis of the Tn-seq results, we identified 42 genes that result in reduced fitness and 45 genes that lead to improved fitness. Within these hits were a number of functional categories that impacted desiccation tolerance.

A major category was that of DNA repair and replication. DNA damage is likely to be one of the most significant stresses of desiccation as many parts of the cell can be remade, but the genome is the template for all necessary cellular components, therefore genome integrity is of the utmost importance. This is supported by the observation that in our screen, disruption of DNA repair and replication genes only resulted in sensitization to desiccation. Of particular interest was the nucleotide excision repair (NER) complex composed of UvrA, UvrB, and UvrC. This complex was highly enriched in our data set based on a Gene Ontology (GO) enrichment for cellular components and we found that deletion of any of these genes resulted in a significant fitness defect. UvrABC is best known to repair thymine dimers that are the result of UV damage, however our desiccations were performed in the dark, making UV an unlikely source of significant DNA damage. UvrABC has also been characterized to repair other DNA lesions, including proteins that have been fused to DNA (40, 41). This may occur as the loss of water results in molecular crowding and loss of hydration shells surrounding proteins and DNA within the cell, causing various cellular components to interact more than they would in a normally hydrated cell (28). Study of DNA damage in desiccated *Bacillus subtilis* spores has previously demonstrated that significant DNA-protein crosslinking occurs during desiccation (42), suggesting that this type of DNA damage likely also occurs in desiccating *S. pneumoniae*.

Single and double stranded breaks have been shown to occur as a result of desiccation (35, 36) and oxidative damage is hypothesized to result from either desiccation or subsequent rehydration of bacteria (43). These other forms of DNA damage would require different repair pathways to resolve specific DNA lesions. When tested, we found that DNA repair pathways which are capable of repairing these types of damage were also required for desiccation tolerance. These pathways include homologous recombination (HR) which would repair double strand breaks and base excision repair (BER) which is capable of repairing modified nucleotides such as oxidatively damaged bases. In addition, we found that nucleotide biosynthesis genes (*prs2, guaA* and *guaB*) involved in maintaining the pool of available nucleotides required for DNA repair and replication also had a decreased fitness in desiccation. Mismatch repair (MMR) was found to have little impact on desiccation tolerance, which can be explained by the fact that MMR generally repairs errors that occur during DNA replication. As we assume the bacteria are metabolically dormant, active DNA replication is unlikely to occur during desiccation. Our finding that deletion of additional DNA repair pathways results in a fitness defect suggests that multiple types of DNA damage are occurring during desiccation and a full complement of DNA repair systems is required for the bacteria to survive after desiccation.

A second category that emerged from our screen was genes that impact structural integrity of the cell. These include genes involved in cell wall and cell division as well as lipid metabolism and envelope biogenesis. During desiccation the volume of the cell decreases while the membrane and cell wall remain their original size (28). This results in dense packing of phospholipids resulting in decreased membrane fluidity and distortion of the membrane which can eventually result in membrane rupture. We found that production of the phospholipid cardiolipin by cardiolipin synthetase (SPD_0185) significantly improves desiccation survival. Cardiolipin is known to increase membrane fluidity which decreases packing of the membrane (44), and thus may be instrumental during desiccation. A structurally sound cell wall also likely helps avoid membrane rupture throughout desiccation as well as during the osmotic shock of rehydration. We found that two class A penicillin binding proteins (PBPs), Pbp1A and Pbp1B, were both important for wild-type levels of desiccation tolerance. The function of these two proteins is still not fully understood, however they are known to be required for maturation of the cell wall as opposed to the construction of nascent peptidoglycan (45). In addition, these genes are synthetically lethal, suggesting they share some functional redundancy in an essential process (46). Loss of type A PBP’s have been characterized to lead to decreased cell-wall stiffness and fewer peptidoglycan crosslinks in *E. coli* and *B. subtilis* (47, 48). Improvements in cell wall integrity by Pbp1A and Pbp2A may increase the bacterium’s resistance to osmotic shock, resulting in improved desiccation survival. It is clear that the condition of the bacterial membrane and cell wall has a large impact on pneumococcal desiccation survival.

Another category of interest from our screen includes metabolic genes and transporters. Previous work has demonstrated that starvation and metal sequestration result in improved desiccation tolerance of *S. pneumoniae* (26). We found multiple sugar transporters, carbohydrate catabolic genes, and a putative metal transporter whose disruption resulted in increased desiccation resistance, which is in agreement with this previous finding. However, the exact mechanism of this improved desiccation tolerance of carbohydrate and metal starved bacteria is unknown. Slower growth could result in smaller cells which will undergo less shrinkage and membrane stress as they desiccate (49). Additionally, slow growth in *Vibrio cholerae* has been shown to improve resistance to osmotic shock (50). More work should be done to understand the impact of decreased growth rate on desiccation tolerance in *S. pneumoniae*.

In order to demonstrate the impact of decreased desiccation tolerance on transmission, the desiccation sensitive mutant Δ*uvrA* was tested in an infant mouse model of transmission. We found that deletion of *uvrA* results in decreased transmission efficiency between mice. It is known that pneumococcal shedding has a large impact on transmission efficiency (51), therefore it was important to demonstrate that Δ*uvrA* did not have a colonization defect that could result in decreased shedding. We found that the bacterial load of Δ*uvrA* in the nasopharynx was the same as wild-type, suggesting colonization density is not the cause of the transmission defect. We suggest that the transmission defect is due to the desiccation sensitivity of our mutant, however we do not have direct evidence that transmission occurs from desiccated bacteria in our murine model. The possibility remains that transmission occurs by direct contact between mice. However, we hypothesize that some of the shed *S. pneumoniae* become desiccated on surfaces in the cage as well as the skin of the pups and the dam. This is supported by the observation that desiccated *S. pneumoniae* remain capable of colonizing a new host (9). Additionally, an association between desiccation tolerance and transmission efficiency has been observed in a pyruvate oxidase mutant, which is both more desiccation resistant and has increased transmission rates in the infant mouse model (26). While these results do not directly demonstrate fomite transmission, they do exhibit a strong correlation between desiccation tolerance and transmission efficiency.

This work has highlighted a number of genetic factors that influence desiccation tolerance of *S. pneumoniae*. In particular, the ability to repair damaged DNA appears to be a key factor that enables bacterial survival and transmission between hosts. Use of DNA damaging agents may be an effective strategy to eliminate bacteria from surfaces. For example, Far-UVC light (222 nm) has been demonstrated to effectively kill infectious bacteria while leaving mammalian skin undamaged (52–54). Utilization of such sterilizing techniques that cause additional DNA damage may prove to be an effective method to decrease the bacterial load on surfaces, thereby reducing pathogen transmission.

## MATERIALS AND METHODS

### Bacterial Strains and Growth Conditions

All experiments were performed with *S. pneumoniae* serotype 2 strain D39 and isogenic mutants, except transmission assays which were performed with serotype 19F strain BHN97 (55). Bacteria were cultivated in a 37°C incubator with 5% CO_2_. Liquid cultures were grown on Todd Hewitt broth (BD Biosciences) supplemented with 5% yeast extract (Fisher Scientific) and 300 U/ml catalase (Worthington Biochemicals) (THY broth). Overnight growth was performed on blood agar (BA) plates which consist of tryptic soy agar (Sigma-Aldrich) with 5% sheep’s blood (Northeast Laboratory Services). Antibiotics were used at the following concentrations: chloramphenicol 4 μg/mL and spectinomycin 200 μg/mL.

### Strain construction

Marked deletion strains were constructed by transforming competent *S. pneumoniae* with PCR products carrying the desired deletion. Allelic exchange PCR products were made using splicing by overlap extension (SOE) PCR as described (56), where the chloramphenicol cassette was spliced to a minimum of 1 kb of sequencing flanking each side of the gene to be deleted. The flanking sequences allow for allelic replacement by double cross-over homologous recombination. Complementation was performed by placing the promoter region, coding sequence, and spectinomycin cassette into a neutral gene locus (SPD_0022). All mutations were confirmed by sanger sequencing or whole genome sequencing. Strains used in this study are listed in table 2.

**Table 2.**
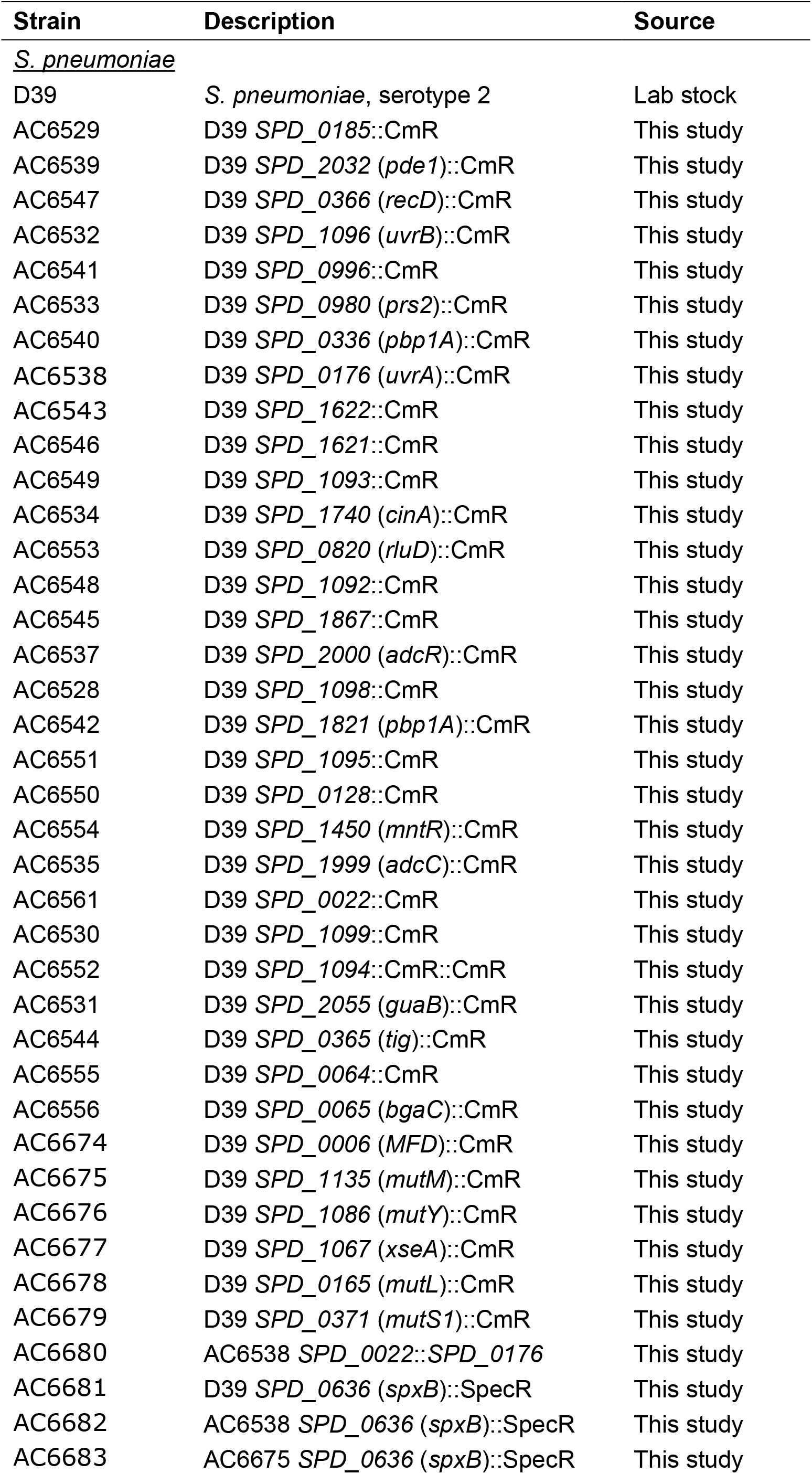
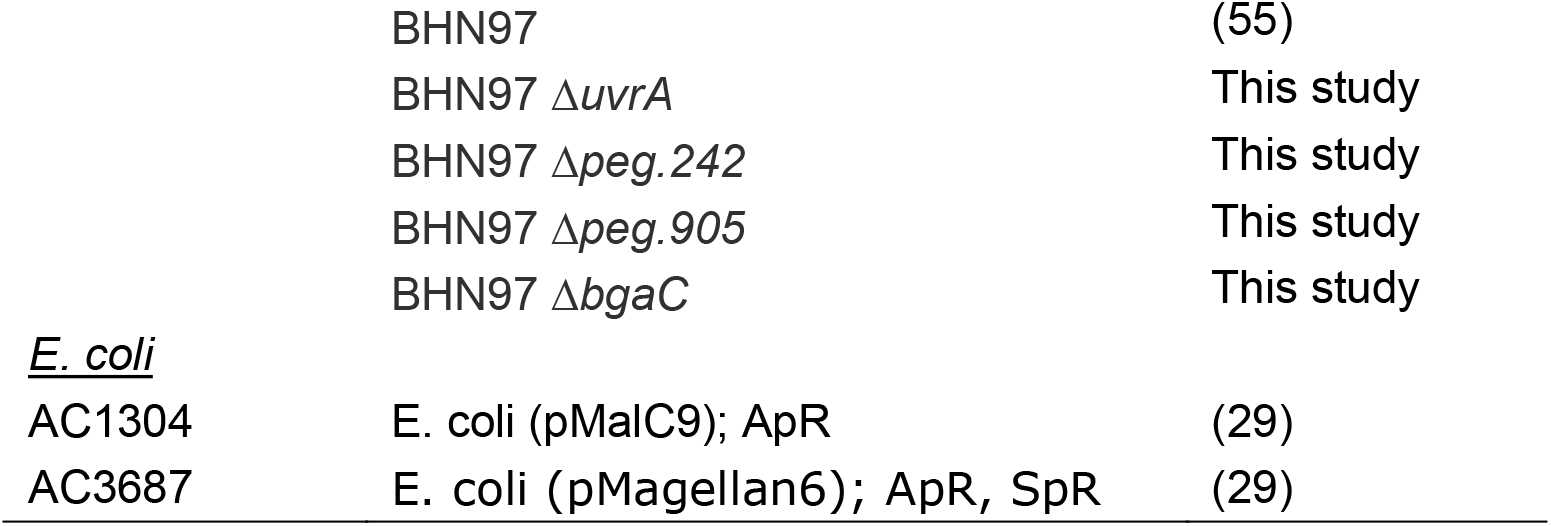
Bacterial strains used in this study.

### Desiccation protocol

*S. pneumoniae* were struck from a frozen glycerol stock onto blood agar plates and grown overnight. Colonies were subsequently resuspended into THY broth and diluted to an optical density at 600 nm (OD600) of 0.1. 50 μL of a 10-fold dilution was then spread onto a blood agar plate and allowed to grow for 16 hours. The resulting semi-confluent colonies were pooled and scraped off a plate using a plastic wedge (Bio-Rad gel releaser 1653320) and split into equal sections. Each section was spread very thinly on a polystyrene petri plate lid using the wedge. Input CFU was quantified by immediate resuspension of the bacteria from several lids in THY and plating of 10-fold dilutions on blood agar. The remaining bacteria were then allowed to desiccate on lids for 2 or 4 days depending on the experiment, after which bacteria were resuspended in THY and plated for viable CFU counts. Bacterial counts were used to calculate percent survival.

Competitions were performed as described above using a 1:1 mixture of unmarked WT and a chloramphenicol resistant mutant to plate the bacterial lawn. Dilutions of bacteria collected from lids on day 0 and day 4 were plated on blood agar and incubated for 16 hours. The resulting colonies were replica-plated onto blood agar containing 4 μg/ml chloramphenicol to assess the ratio of mutant to WT. All were done with 5 to 10 biological replicates.

### Transposon Library Construction

The transposon library was constructed as previously described (57). Briefly, in vitro transposition was performed using purified transposase MarC9, genomic DNA, and the mini transposon magellan6, which contains a spectinomycin-resistance gene. Transposed DNA was then transformed into competent *S. pneumoniae*, and bacteria carrying a transposon were selected for by plating on blood agar supplemented with 200 μg/ml spectinomycin. This pool of mutants was grown up in THY, then collected and frozen down in 20% glycerol (final concentration) for further experimentation.

### Desiccation Tn-seq screen

Libraries that were previously frozen were plated on blood agar and grown for 16 hours. Each biological replicate consisted of ten 150 mm diameter blood agar plates. The following day the bacterial lawns were collected, mixed together, and desiccated as described above. Three input samples were collected immediately after spreading on lids and plated on blood agar. Five output samples were collected after 2 days of desiccation and plated on blood agar. After overnight growth, bacteria were collected and frozen as glycerol stocks for future isolation of genomic DNA.

### Sequencing and Analysis Pipeline

Genomic DNA was isolated using the DNeasy Blood and Tissue Kit (Qiagen, 69504). Samples were prepared for sequencing using the HTML PCR method (29). Briefly, genomic DNA was sheared via sonication in a cuphorn sonicator and poly-C tails were added to the 3’ ends of all fragments using terminal deoxynucleotidyl transferase. Transposon junctions were amplified through PCR amplification using primers specific for the Magellan6 transposon and the poly-C-tail. A subsequent nested PCR was performed to add unique barcodes to each sample. Sequencing was performed as 50 bp single-end reads on an Illumina HiSeq 2500 at the Tufts University Core Facility.

Fitness was calculated as previously described (57). Briefly, reads were mapped to the D39 genome using Bowtie (58). Transposon insertions at each gene locus were quantified for all input and output samples and the data were normalized to the total number of reads in each sample to account for slight variations in read depth. Fitness for each unique insertion was calculated as previously described (29). No change is quantified as a fitness of 1, representing a neutral gene. Increased presence in the output results in a fitness greater than 1, while decreased presence in the output produces a fitness less than 1. Fitness values were then normalized against a list of neutral genes from D39 to artificially set those gene’s fitness to 1 and the same factor was used to normalize all other fitness values. Mean fitness of a gene was calculated by averaging all unique insertions across a gene. A minimum cutoff of 4 unique transposon insertions per gene was applied in addition to a read cutoff of 15 reads per transposon insertion. Next a fitness cutoff was applied to remove all genes with less than a 20% fitness change from the neutral fitness of 1. Finally, statistical significance was determined using a sample t-test with Bonferroni correction for multiple comparisons.

### UV irradiation challenge

Strains were grown up in THY broth to mid-log phase. Cultures were washed and resuspended in PBS, then 50 μl was spotted onto parafilm. Bacteria were exposed to 15 millijoules of ultraviolet light (254 nm) using a Stratagene UV crosslinker. Bacteria from before and after UV treatment were plated on blood agar to quantify CFU and this was used to calculate percent survival. All were done with six biological replicates over two separate days.

### Transmission assay

This assay was performed as previously described (26). Briefly, litters of 4-day old C57/BL6 infant mice were split into two equal groups. The first group was colonized with either wild-type or mutant serotype 19F strain BHN97, termed the donor mice. The other group was left uncolonized and referred to as the contact mice. All mice from the litter were then placed back in the cage with the dam. Transmission was tracked over the course of 10 days by tapping the nares of the contact mice against a blood agar plate. Detection of bacteria on two subsequent days was defined as a transmission event. At the conclusion of the transmission experiment, all mice were sacrificed, and the level of nasopharyngeal colonization was quantified to ensure that varied transmission levels are not the result of increased or decreased shedding from the donor mice.

## ACKNOWLEDGEMENTS

This research was supported by National Institutes of Health training grant GM139772 (A.J.M.).

